# Stabilization demands of walking modulate the vestibular contributions to gait

**DOI:** 10.1101/2020.09.30.319434

**Authors:** Rina M. Magnani, Sjoerd M. Bruijn, Jaap H. van Dieën, Patrick A. Forbes

## Abstract

Stable walking relies critically on motor responses to signals of head motion provided by the vestibular system, which are phase-dependent and modulated differently within each muscle. It is unclear, however, whether these vestibular contributions also vary according to the stability of the walking task. Here we investigate how vestibular signals influence muscles relevant for gait stability (medial gastrocnemius, gluteus medius and erector spinae) – as well as their net effect on ground reaction forces – while humans walked normally, with mediolateral stabilization, wide and narrow steps. We estimated local dynamic stability of trunk kinematics together with coherence of electrical vestibular stimulation (EVS) with muscle activity and mediolateral ground reaction forces. Walking with external stabilization increased local dynamic stability and decreased coherence between EVS and all muscles/forces compared to normal walking. Wide-base walking also decreased vestibulomotor coherence, though local dynamic stability did not differ. Conversely, narrow-base walking increased local dynamic stability, but produced muscle-specific increases and decreases in coherence that resulted in a net increase in vestibulomotor coherence with ground reaction forces. Overall, our results show that while vestibular contributions may vary with gait stability, they more critically depend on the stabilization demands (i.e. control effort) needed to maintain a stable walking pattern.

## Introduction

The stability of walking is commonly assessed as the ability to maintain upright locomotion in the presence of self-generated and/or external perturbations ^1-3^. To ensure stable walking, the nervous system relies on the integration and modulation of sensory signals from visual, somatosensory and vestibular sources to generate ongoing and/or corrective postural responses throughout the different phases of the gait cycle ^4-7^. The vestibular system, for instance, which encodes signals of head movement in space ^8^, is assumed to contribute to gait stability because impaired gait is often observed in vestibulopathic patients ^9,10^. Recent studies have further revealed that vestibular contributions to locomotion undergo phase- and muscle-specific responses that appear to align with each muscle’s functional role in gait stability ^11,12^. Specifically, the vestibular contributions to mediolateral stability during walking may be related to mediolateral foot placement produced by muscles around the hip ^11,13,14^ and to ankle torque at push-off driven by muscles of the ankle ^15^. In the sagittal plane, however, corrective responses to a vestibular disturbance are almost entirely absent throughout the gait cycle ^16^. Because stability in the sagittal plane is maintained largely passively ^17^ – due to passive dynamics of the legs ^18^ – it may be possible that the control of whole-body stability during locomotion requires less feedback-driven control as compared to the frontal plane ^16^. Thus, the question arises whether changes in the stability of walking in the mediolateral plane also influences the vestibular contribution to stability during locomotion. Therefore, our aim is to identify how vestibular contributions to balance are modulated across varying stabilization demands of walking by characterizing changes in coupling of vestibular input to motor outputs during externally-imposed and natural variations in stability.

Experimentally, gait stability can be improved by adding external lateral stabilization to the body, thereby removing the need for the nervous system to control upright balance in the mediolateral direction ^19-21^. Under these stabilized conditions, step width variability, trunk and pelvis motion and premotor cortical involvement all decrease compared to normal walking ^22-25^. These observations indicate a reduced need to actively control gait stability, and as a result, may also diminish the necessity for vestibular sensory feedback control. This may be similar to observations during upright standing, where compensatory responses to mediolaterally-directed vestibular disturbances are absent when participants are stabilized in the mediolateral direction, despite having to maintain balance in an anterior-posterior direction ^26^. Therefore, we first hypothesized that external lateral stabilization of the body would diminish the muscle and whole-body responses to a mediolateral vestibular disturbance during locomotion.

Natural changes in gait are also thought to influence stability. For instance, adopting a wider step width has been described as a response to decreased lateral stability during locomotion ^22,27^, and seems to be an effective approach to increase margins of stability (i.e. base of support) in older adults ^28,29^. If the vestibular system influence on gait is tightly coupled to stability, then similar to external stabilization, we expected that the evoked muscle and whole-body responses to a vestibular disturbance should diminish when walking with wider steps. On the other hand, narrow-base walking requires increased effort to control gait stability in the frontal plane in young and old adults ^30,31^, since the margins of stability, or tolerance for errors, are substantially reduced ^32,33^. These observations, however, seem to contrast with measures of local dynamic stability, which quantify the likelihood of departing from a steady-state gait pattern in the absence of external disturbances, and instead indicate increased gait stability during narrow-base walking and decreased gait stability during wide-base walking ^31,34^. We aim to test a potential explanation for this conflict by examining whether the increased local dynamic stability observed when walking with narrow step width is (partially) subserved by increased vestibular sensory feedback. Indeed, support for this is seen by the additional contribution of vestibular signals (via the lateral vestibular nuclei) to limb muscle activity in mice when walking on a narrow beam that is absent when walking on level ground ^35^. Here, we tested these hypotheses by comparing muscle and whole-body responses evoked by a vestibular disturbance – together with estimates of stability – across normal, mediolaterally stabilized, wide-base and narrow-base walking.

## Methods

### Participants

We measured 23 healthy young adults between 24 and 33 years recruited from the university campus. Twelve participants were excluded during data analysis due to technical problems in the collection of electromyography (n=3) or kinematic data (n=9) in any of the trials recorded. Here we present results from eleven young adults (four females, 28.5±2.9 years old, 71.6±8.6 kg and 1.77±0.10 m, body mass index 22.2±3.1 kg/m^2^). Exclusion criteria included self-reported history of injury and/or disfunction of the nervous, musculoskeletal or vestibular systems, or the use of medications that can cause dizziness. Participants were also instructed not to participate in intense physical exercise on the day of the experiment. The participants agreed to participate in the study by signing the informed consent form, and the study was approved by the VU Amsterdam Research Ethics Committee (VCWE-2017-158).

### Electrical vestibular stimulation

A continuous electrical vestibular stimulus (EVS) was used to deliver an isolated vestibular disturbance to participants during all walking trials. Coupling of the electrical stimulus with muscle activity and ground reaction forces was quantified over the gait cycle to determine the magnitude and timing of the vestibular contribution to ongoing muscle and whole-body responses. The electrical stimulus modulates the afferent firing rate of both semicircular canal and otolith afferents ^36-38^, and when delivered in a binaural-bipolar configuration, EVS evokes a sensation of head roll rotational velocity ^39^ about an axis directed posteriorly and superiorly by 18° relative to the Reid plane ^40-42^. When the head is facing forward, this stimulus configuration results in a postural response in the frontal plane to compensate for the induced roll error signal ^16,26,43-46^.

The electrical stimulus was applied to participants using flexible carbon rubber electrodes (9 cm^2^). The electrodes were coated with Spectra 360 electrode gel (Parker La, USA) and fixed to participants’ mastoid processes using adhesive tape and an elastic head band. The stimulus was delivered as an analog signal via a data acquisition board (National Instruments Corp., Austin, TX, USA) to an isolated constant current stimulator (STMISOLA, Biopac, Goleta, CA, USA). All participants were exposed to the same stochastic EVS designed with a limited bandwidth of 0 to 25 Hz ^47^, zero-mean low-pass filtered white noise, 25 Hz cutoff, zero lag, fourth-order Butterworth, peak amplitude of 5.0 mA, root mean square of ∼ 1.2 mA, lasting 8 minutes and created with Matlab software (MathWorks, Natick, MA, USA). Because this binaural-bipolar stimulus oscillates around a zero-mean, the imposed sensations of roll motion, and the accompanying compensatory responses, occur in both a left and right direction.

### Protocol

Participants walked on a dual-belt treadmill at a belt speed of 0.8 m/s in four different conditions: normal walking, stabilized walking, wide-base walking, and narrow-base walking. During normal walking and stabilized walking, participants were instructed to walk with their naturally preferred step width. Stabilized walking was achieved using a custom-made spring-loaded mediolateral pelvic stabilization frame. The stabilization frame was attached through springs to two carts that allowed for movement in the anterior-posterior direction. The springs were pre-tensioned to provide a stabilizing stiffness of 1260 N/m ^48^ and the height of the carts was aligned with the height of the pelvis for each subject ^49^. During wide-base walking, participants were instructed to increase their step width beyond the approximate width of their hips. During narrow-base walking, participants were instructed to adopt a step width smaller than both the width of their hips and their usual step width. Throughout wide- and narrow-base trials, participants received repeated verbal instruction to maintain wider and narrower step widths, respectively, compared to normal walking. Participants walked in each condition for 8 minutes while being exposed to continuous EVS and were guided by the beat of a metronome at 78 steps/min to control for effects of varying cadence on vestibular contributions during locomotion ^11,12^. The walking speed of 0.8 m/s and cadence 78 steps/min were chosen to replicate the conditions of Dakin et al. (2013) ^11^. These walking parameters also ensure that vestibular-evoked balance responses, which are known to decrease as velocity and cadence increase ^11,12^, could be measured throughout the gait cycle.

Prior to starting the experiments, participants were allowed to walk for 3-4 minutes to familiarize themselves with walking on the treadmill at the specific cadence and with the electrical stimulus. In addition, participants walked for 2 minutes in each condition before the electrical stimulus was applied, which together with the familiarization period was considered a sufficient exposure period to remove acclimatization effects of walking on a treadmill ^50,51^. Trial order for the different walking conditions was also randomized for each subject and subjects were given a short 5 minute break between trials to limit the influence of any long-term habituation to the electrical stimulus throughout the walking trials ^52^. Finally, participants maintained their head in a slightly extended position with the Reid’s plane pitched ∼18° up from horizontal ^40,41^ by keeping a headgear-mounted laser on a target located 3 m in front of them. This head position was chosen to maximize the amplitude of vestibulomotor balance responses in the mediolateral direction ^41,53,54^.

### Instrumentation

Kinematic data were recorded using a 3D motion capture system (Optotrak, Northern Digital Inc., Waterloo, Ontario, Canada) sampling at 100 samples/s. Clusters of three light emitting diodes (LED) were positioned at the occipital lobe, the spinous process of the sixth thoracic vertebra (T6), the posterior superior iliac spine and at the calcaneus bilaterally. Ground reaction forces (GRF) were measured from each belt by force plates embedded in the treadmill (Motekforce Link, The Netherlands) at a sampling rate of 200 samples/s. In addition to analysis of the coupling between EVS and mediolateral forces as described below, these signals were used for the identification of toe-off and heel-strike and gait events.

Surface electromyography (EMG) (TMSI Porti system, TMSI Enschede, the Netherlands) was collected at 2000 samples/s bilaterally from the medial gastrocnemius, gluteus medius and erector spinae muscles, using pairs of disposable self-adhesive Ag/AgCl surface electrodes (Ambu, Balerrup, Denmark; model Blue sensor; diameter 30×22mm) for each muscle. Electrodes were placed over the recorded muscles according to SENIAM electrodes placement recommendations ^55^ after abrading and cleaning the skin with alcohol. A reference electrode was placed over the medial bony part of the left wrist (styloid process). The three muscles measured were chosen based on their supposed roles in different stabilizing strategies that may be employed throughout the gait cycle and across walking conditions. The medial gastrocnemius muscle (and other ankle plantarflexors) act as the foot’s prime movers during normal walking ^56^ and are most sensitive to vestibular input in the late stance phase ^11,12,15^ when modulating push-off force. The gluteus medius muscle contributes to foot placement strategies; its activity is correlated to the next foot placement ^57,58^ and is most sensitive to imposed vestibular errors just prior to heel strike ^11^. Finally, trunk muscles serve to directly influence trunk motion relative to the pelvis and are primarily activated to stabilize the trunk during weight transfer around heel strike ^59,60^. Therefore, erector spinae muscle may be especially suited to contribute to angular momentum control of the torso during narrow base walking ^61^ when push-off modulation and foot-placement are rendered ineffective.

### Data analysis

Force-plate data were used first to calculate center of pressure positions, which were in turn used to identify heel contacts ^62^. From these estimates, stride time was calculated as the duration between two consecutive heel strikes of the same foot and the step width was determined as the mediolateral distance between the centroids of the feet cluster markers at heel strike. These measures of limb kinematics (stride time and step width) were compared across walking conditions to determine whether participants adhered to our instructions (that is, walk at the metronome-guided cadence and with a modified step width). We compared muscle activity across each walking condition after rectifying and low-pass filtering (20 Hz cutoff, zero lag, sixth order Butterworth) the EMG signals and then time-normalizing the data per stride and averaging the data across the 256 strides. Prior to averaging, each rectified and low-pass filtered EMG signal was normalized to the maximum amplitude across all conditions.

To assess changes in local dynamic stability during the different walking conditions, we estimated the local divergence exponent (LDE). Measures of local dynamic stability for walking indicate the ability for a participant to return to the steady-state periodic motion after infinitesimally small perturbations ^63^, which occur, for example, through natural variability in the walking surface or the neuromuscular system. These measures have been shown to be particularly useful for detecting patients at risk of falling ^3^. The LDE measures the exponential rate of divergence of neighboring trajectories of a state space constructed from kinematic data of gait ^64^, whereby an increasing LDE indicates reduced stability. We calculated the LDE using Rosenstein’s algorithm ^65^ and as input the velocity of a maker placed over the T6 vertebrae, which was estimated using a three-point differentiation of the position trace ^66^. Velocity time series were first resampled so that each time series of 256 strides (the minimum number of strides collected from every participant) contained 25600 samples. The LDE was then calculated as the slope of the mean divergence curve, whose horizontal axis was normalized by stride time from 0-0.5 stride ^67,68^.

To examine the vestibulomotor coupling between the input stimulus (i.e. EVS) and motor output (EMG and ground reaction forces) across our conditions, we computed the time-frequency coherence and gain assuming a linear stimulus-response relationship ^69^. Prior to estimating coherence and gain, EVS, EMG and ground reaction forces were cut into segments synchronized to heel strikes. Based on the symmetry of our walking conditions, we chose to align the data on each limb’s heel strike in order to pool responses from muscles in the left and right limbs. Symmetry of the evoked muscle responses was confirmed prior to pooling the data (see Statistical analysis). This approach, however, was not possible for coherence and gain between EVS and GRFs since in our narrow-base walking condition because participants regularly made contact of one limb with the opposing belt such that forces from each limb could not be measured on the separate force plates. Therefore, forces from both plates were first summed before estimating the coherence. To avoid distortion in the coherence estimates at the beginning and end of the signal, each stride was padded with data from the neighboring strides (50%). EVS and rectified EMG were low-pass filtered (100 Hz cutoff, zero lag, sixth-order Butterworth) and down-sampled to 200 samples/s. To account for stride- to-stride variation, stride duration was normalized by resampling the data according to the average stride duration of all trials. This normalization was performed on the auto-spectra of the EVS, EMG and force signals, as well as on their cross-spectra (see below).

Our analysis of coherence and gain was performed based on continuous Morlet wavelet decomposition ^15,70^ using equations (1) and (2):

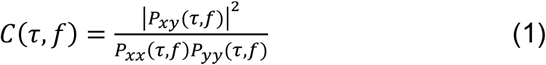

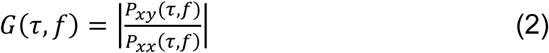

where *Pxy*(*τ, f*) (*τ* and *f* denote the stride time and frequency, respectively) is the time-dependent cross-spectrum between the EVS and rectified EMG or GRF, and *Pxx*(*τ, f*) and *Pyy*(*τ, f*) are the time-dependent auto-spectra of the EVS and rectified EMG or GRF, respectively. Coherence ranges from zero to one, and provides a measure of the linear relationship between two signals ^71^ as well as an estimate of their coupling (i.e. shared variance) at each frequency ^72^. Because coherence is normalized to the auto-spectra of the input and output signals, it can be sensitive to variation in non-vestibular contributions to the measured output signal (i.e. magnitude changes of EMG or ground reaction forces). Gain on the other hand indicates the magnitude of the output relative to the input and is not normalized by the output signal power spectrum; as a result, it is not expected to change even if non-vestibular input leads to changes in output signal magnitude. A preliminary evaluation of our results indicated that similar to previous studies ^12,15^, both coherence and gain followed parallel changes in magnitude and timing across the four walking conditions. This suggests that the modulation in coherence was not simply dependent upon the magnitude of the measured output signals, and as a result, we present only the coherence to describe the modulation of vestibular contributions to mediolateral stability. Finally, we evaluated the coherence between EVS and both EMG and GRFs to estimate muscle-specific vestibular contributions and to provide a net estimate of the vestibular input to ongoing locomotor behavior, respectively.

### Statistical analysis

We compared all gait parameters (stride time, step width and local dynamic stability) between the four conditions using one-way repeated measures ANOVAs. Subsequently, we performed planned pairwise comparisons (t-test) between normal walking condition and each modified walking condition (i.e. normal vs. stabilized; normal vs. narrow and normal vs. wide-base walking). EVS-EMG coherence and EVS-GRF coherence were used to identify the phase-dependent coupling between vestibular stimulation and motor responses throughout the walking cycle. For each participant, coherence was defined as significant for those points in the gait cycle where it exceeded 0.018, corresponding to p<0.01 (for 256 strides) in view of the bi-dimensional nature of the correlations ^15^. To determine whether our walking manipulations modified the EVS-EMG coherence when compared to normal walking, we performed cluster-based permutation tests (paired t-tests, 5000 permutations) ^73^ between conditions aimed to identify whether the time-frequency-coherence spectra significantly differed from the normal condition. In doing so, we did not disregard non-significant coherence values. Since our initial analysis showed no significant between leg differences in coherence, we averaged coherence values over legs.

## Results

### Effects of condition on gait parameters, muscle activity and local dynamic stability

To characterize changes in gait across walking trials, we first evaluated the gait parameters, muscle activity and stability measures. During all trials, participants were able to maintain stable upright locomotion while exposed to the stochastic electrical stimulation. Despite walking to the beat of a metronome in all conditions, we found a significant main effect of condition on stride time (F(3,30)=4.14; p=0.014; η^2^=0.127). Pairwise comparisons revealed that stride time increased by approximately 6.12±15.65% (0.09±0.02 s; p=0.018) during stabilized walking when compared to normal walking (Figure 1a), but it did not change significantly during either wide-base (p=0.928) or narrow-base walking (p=0.155). As intended, walking condition significantly affected step width (F(3,30)=73.4; p<0.001; η^2^=0.786, see Figure 1b). Consistent with previous results ^20,23-25^, pairwise analysis revealed that participants reduced their step width by 65.31±40% (0.18±0.02 m; p<0.001) during stabilized walking, in spite of no explicit instruction to do so. We also found that participants adhered to the step width instructions during the other two conditions, increasing step width by 30.25±20% (0.08±0.02 m; p<0.001) during wide-base walking and reducing step width by 38.74±5% (0.11±0.01 m; p<0.001) during narrow-base walking (Figure 1b).

**Figure 1:**
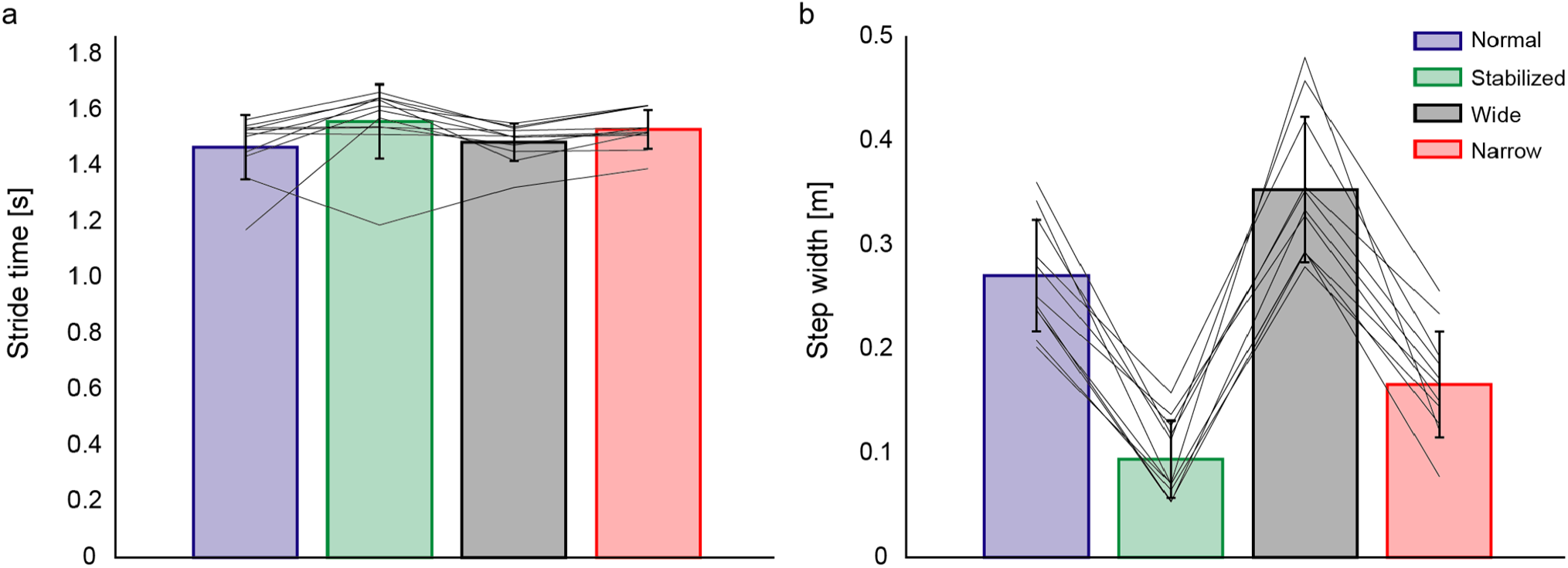
Spatiotemporal parameters of gait during normal walking (blue), walking with external lateral stabilization (green), wide-base walking (black) and narrow-base walking (red). Mean values (bars), standard deviation (error bar) and participants individual changes across conditions (lines) for the stride time (a) and stride width (b) for the four conditions.

As expected, when compared to normal walking, significant differences in mediolateral ground reaction forces ^30,31,74^ during all other walking conditions were observed primarily during single-support phases of the gait cycle (see Figure 2a). Forces decreased when subjects were externally supported or walked with a narrow-base, and increased when subjects walked with a wide-base (see Supplementary Figure S1). In the medial gastrocnemius and gluteus medius muscles, peak activity in each muscle occurred during the double stance phase (∼50% of the stride cycle) and after heel strike (∼20% of stride cycle), respectively. EMG responses for these two muscles were for the most part overlapping in all conditions, with only limited significant differences observed during the stance phase for some of the conditions relative to normal walking (see Figure 2b/c and Supplementary Figure S1). Muscle activity in the erector spinae demonstrated more complex phasic activity throughout the gait cycle with two peaks occurring just after heel strike of each limb. Limited significant differences were observed during stabilized walking and narrow-base walking when compared to normal walking (see Figure 2d and Supplementary Figure S1).

**Figure 2:**
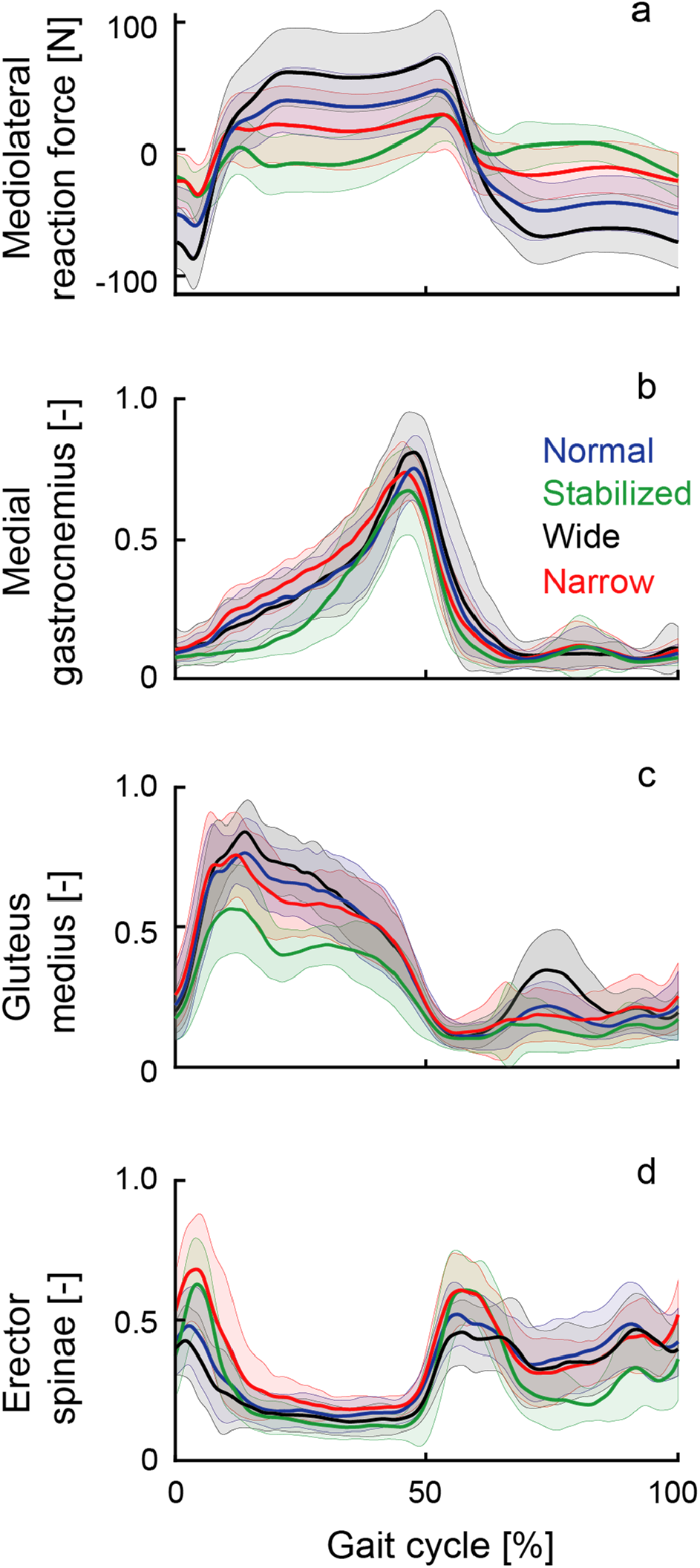
Ground reaction force and electromyographic amplitudes throughout the gait cycle during normal walking (blue line), walking with external lateral stabilization (green line), wide-base walking (black line) and narrow-base walking (red line). Mean values (lines) and standard deviation (shadow area) of the ground reaction force (a) and the EMG envelopes of the medial gastrocnemius (b), gluteus medius (c) and erector spinae (d) muscles for all conditions. For each muscle, EMG signals were normalized to the maximum amplitude across all conditions.

Our manipulations also had a significant effect on mediolateral stability, as expressed by the local divergence exponent (F(3,30)=129; p<0.001; η^2^=0.712). Pairwise comparisons showed that participants had a higher stability (i.e., lower LDE values) during the stabilized (p<0.001) and narrow-base walking conditions (p=0.019) compared to normal walking. Although the mean local divergence exponent was highest (i.e. lowest stability) during wide-base walking, this was not significantly different from normal walking (p=0.265) (Figure 3).

**Figure 3:**
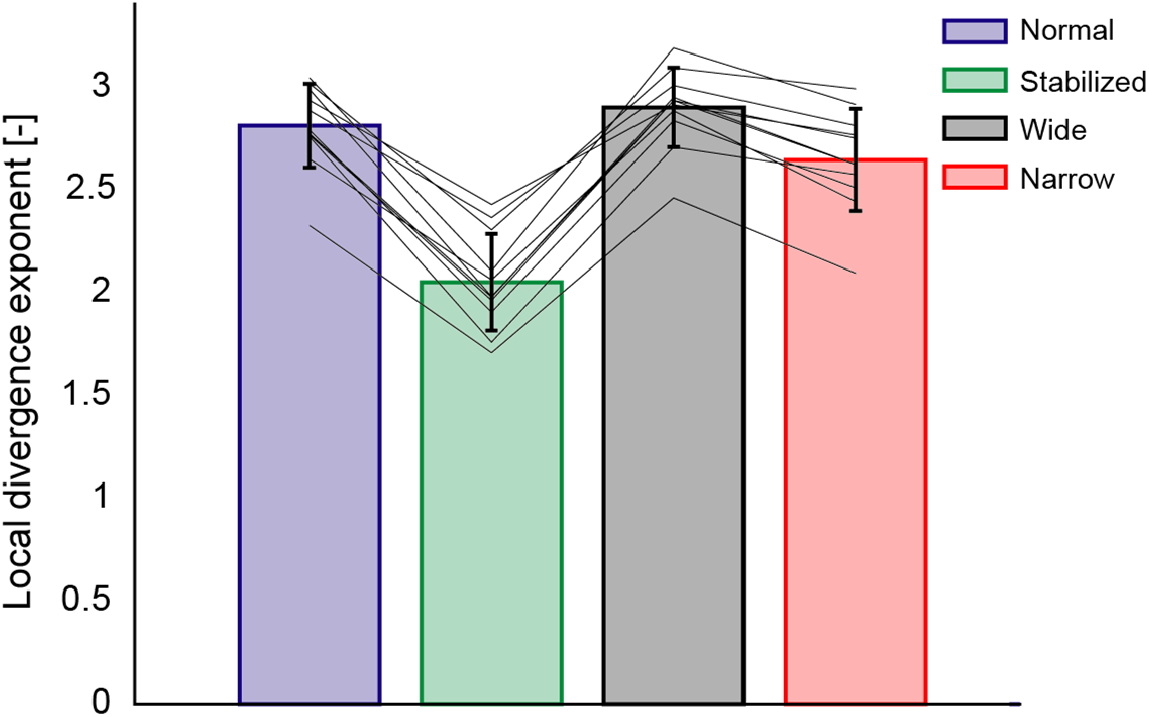
Local divergence exponents during normal walking (blue), walking with external stabilization (green), wide-base walking (black) and narrow-base walking (red). Mean values (bars), standard deviation (error bar) and participants individual changes across conditions (lines) for the evaluated conditions.

### EVS-mediolateral GRF and EVS-EMG coherence in normal walking

We next characterized the coupling of the electrical stimulus with both the ground reaction forces and muscle activity in normal walking as a baseline for comparison to our manipulated conditions (Figure 4 – first column). During normal walking, significant EVS-GRF coherence was seen in all participants over the entire gait cycle, with phase-dependent group mean responses that peaked during single stance (Figure 4 top row). Significant phase-dependent EVS-EMG coupling was also prominent in the mean responses during normal walking, but muscle-specific variations were observed: coherence peaked in mid stance in the medial gastrocnemius, just before heel strike in gluteus medius, and at heel strike and at mid stride (though at a lower magnitude) for the erector spinae. Consistent with previous reports ^11,12,15^, peak coherences did not align with peak EMG for any of the muscles, further confirming that vestibular contributions do not depend purely on the excitation of the motoneuron pool. In addition, as commonly observed in vestibular-evoked muscles responses during standing ^75^, the bandwidth of significant EVS-EMG coherence spanned ∼0-25 Hz while significant EVS-GRF coherence was observed from ∼0-10 Hz (Figure 4).

**Figure 4:**
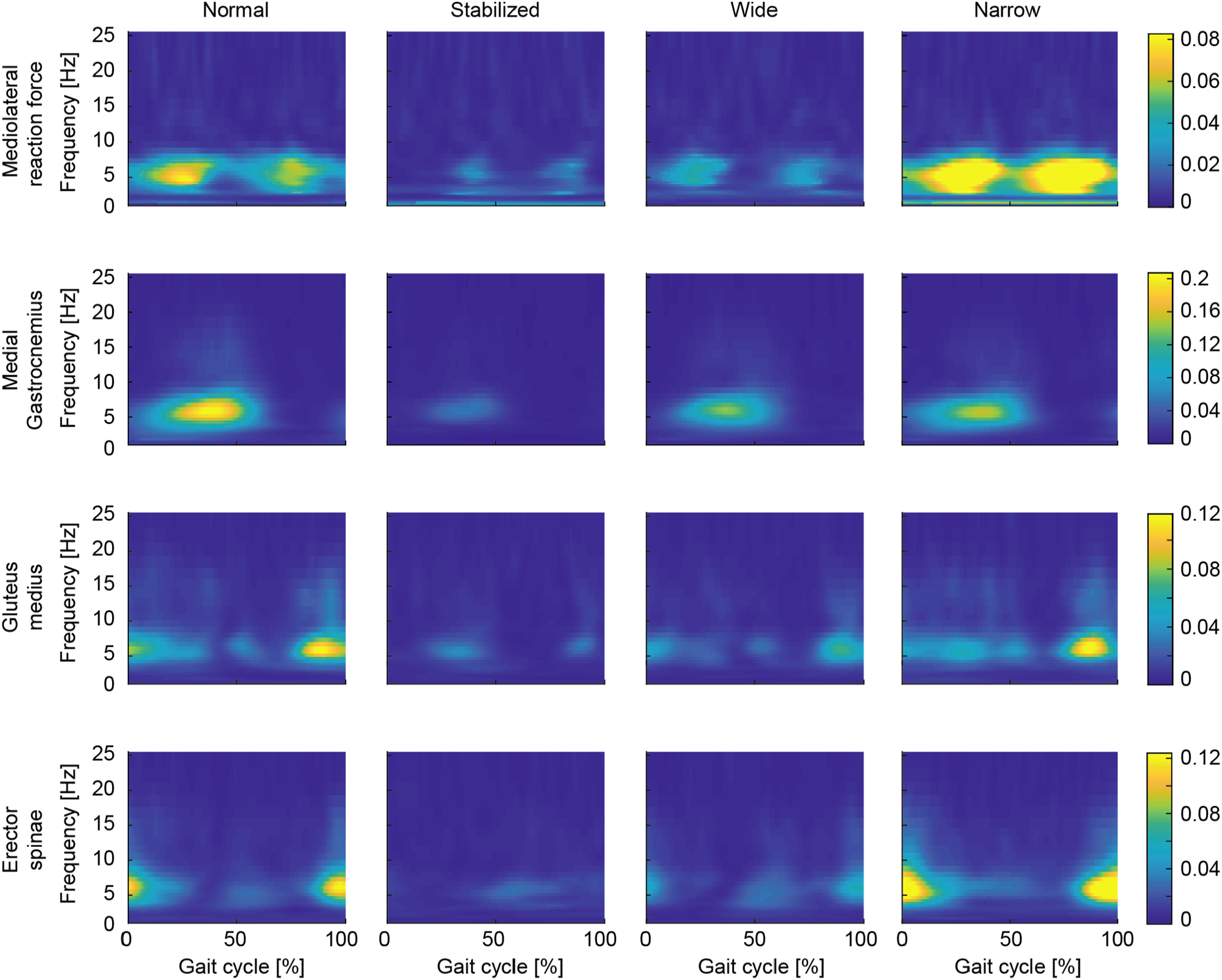
Coherence plots of EVS-GRF (first row) and EVS-EMG for medial gastrocnemius (second row), gluteus medius (third row) and erector spinae (fourth row) for normal walking (first column), walking with external stabilization (second column), wide-base walking (third column) and narrow-base walking (fourth column). Coherence magnitude is indicated by the color bars.

### Stabilization demands but not dynamic stability modulate EVS-GRF and EMG-EVS coupling

To establish the effects of stabilization demands on vestibulomotor coupling, we examined the difference in coherence between normal walking and all other conditions (Figure 5). During externally stabilized walking, both EVS-GRF and EVS-EMG coherences decreased significantly relative to normal walking (see Figure 4 and 5). More specifically, the reduced coupling during the stabilized condition was primarily observed during the periods of peak coherence in normal walking (i.e. single stance for GRF and medial gastrocnemius, and before/at heel strike for gluteus medius and erector spinae). Although the increased stride time (i.e. decreased cadence) during stabilized walking (see Figure 1) may have acted as a confounding factor to these changes ^11^, this effect commonly increases vestibulomotor responses in contrast to the observed decrease in coherence seen here. During wide-base walking, EVS-GRF and EVS-EMG coherences were also significantly decreased compared to normal walking (Figure 5). The EVS-GRF coherence decreased over the majority of the gait cycle with the most prominent changes observed during the periods of peak coherence seen in the normal condition. Further, EVS-EMG coherence was reduced for all three muscles in wide-base walking, again with the greatest differences observed at instants of peak coherence in normal walking (Figure 5). Taken together, the results of stabilized and wide-base walking show a reduction in vestibular input to the net muscle activity of the body (i.e. GRFs), which is driven at least in part by the three muscles measured when stabilization demands (but not dynamic stability) are decreased.

**Figure 5:**
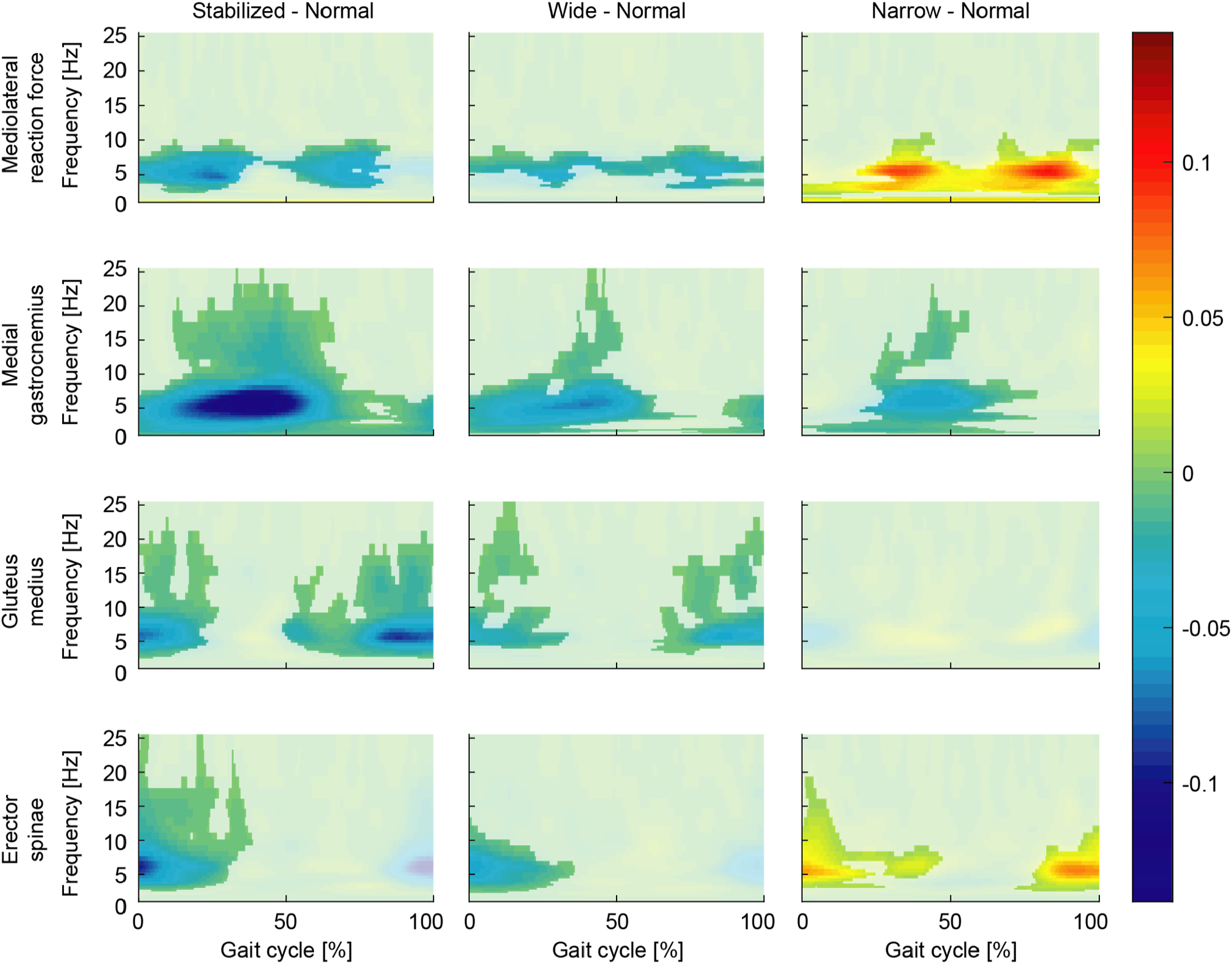
Differences in time-frequency coherence between the normal walking condition and walking with external lateral stabilization (first column), wide-base walking (second column) and narrow-base walking (third column). Color coding refers to the difference in coherence between two walking conditions (e.g. normal minus stabilized) for the EVS-GRF coherence (first row) and EVS-EMG coherence of medial gastrocnemius (second row), gluteus medius (third row) and erector spinae (forth row) muscles. For illustrative purposes, differences that were not significant were plotted slightly opaque.

During narrow-base walking, we observed more complex changes in coupling between EVS and GRF and between EVS and muscle activity. Figures 4 and 5 together show that EVS-GRF coherence during narrow-base walking significantly increased compared to normal walking over the entire gait cycle. While this was matched by an increase in EVS-EMG coherence in the erector spinae muscle, we found unchanged EVS-EMG coupling in the gluteus medius muscle and a decrease in coherence in the medial gastrocnemius muscle. These more complex changes in vestibular-evoked motor responses suggest that while the net output of the vestibular-evoked muscle activity (i.e. EVS-GRF coherence) increases with increased stabilization demands, as well as the dynamic stability (see Figure 3), this trend is not reflected in EVS-EMG coupling of all muscles.

## Discussion

We characterized how coupling of vestibular input with muscle activity and ground reaction forces modulate as a function of the stabilization demands during locomotion. We found that as participants walked with decreased stabilization demands through either external stabilization or wider step widths, coherence between electrical vestibular stimulation and both muscle activity and ground reaction forces decreased compared to normal walking. These overall reductions in vestibulomotor coupling were accompanied by an increase or no change in the stability of the gait pattern – measured as a decreased or constant local divergence exponent – during stabilized and wide-base walking, respectively. In contrast, the increased stabilization demands of walking with narrow steps invoked complex changes in vestibulo-muscular coupling that increased or decreased specific to each muscle’s involvement in correcting for the imposed vestibular error. Nevertheless, these changes in vestibulo-muscular coupling increased the collective contribution of vestibular signals to the ground reaction forces and occurred together with a decrease in the local divergence exponent (i.e. increased gait stability). This suggests that participants maintained a more stable gait pattern during narrow walking that was at least partially subserved through increased use of vestibular feedback. Ultimately, these results indicate that vestibular contributions to gait stability may be modulated with frontal plane stability, but that they more specifically depend on the stabilization demands (i.e. control effort) required to maintain a stable gait pattern and not the stability of the gait pattern itself.

When participants walked with external stabilization, the stability of the gait pattern increased (i.e. decreasing LDE) while vestibular-evoked muscle and force responses decreased as compared to normal walking. Both of these results are not entirely surprising since the control of mediolateral motion is aided by the forces generated by the springs ^19,20,25^. As a result, there is a reduced reliance on vestibular signals to maintain upright locomotion during stabilized walking. This is similar to the task dependent reductions in vestibular input observed during standing ^44,45,76-78^ when participants are externally supported; stimulus-evoked responses are suppressed since the vestibular feedback is no longer relevant to balancing the body. Our results reveal that these task dependent changes in the vestibular control of standing also apply during the more dynamic task of walking. In addition, they also support the proposal that anteroposterior control of whole-body stability during locomotion is controlled passively ^79^. By making the body passively stable in the mediolateral direction, we saw a reduction in vestibular-evoked response that matched the near absence of vestibular contributions when the vestibular error is directed in the anterior-posterior direction ^16^.

When participants walked with a wide base, vestibular-evoked muscle and force responses also decreased in a manner that parallels the effects of wide stance during standing ^44,80^. Walking (and standing) with a wide foot placement increases the base of support, and the passive stiffness in the frontal plane. The current results show that the corrective contribution of vestibular signals during walking with a wide base decrease in a manner similar to the effects seen during external stabilization. Our measure of dynamic stability, however, did not follow the same trend. Instead, we observed a slight (albeit non-significant) increase in the local divergence exponent compared to normal walking. This aligns with previous estimates of a constant or decreased dynamic gait stability during wide-base walking ^30,34^. A key difference between stabilized and wide-base walking is that in the former, increased gait stability and upright balance is an inevitable result of the external support. Wide-base walking, on the other hand, despite the increased base of support, still demands active stabilization, and the manner in which this is achieved differs from normal walking. For example, push-off modulation and step-by-step foot placement precision both decrease when walking with wide steps ^30^. In addition, the increased moment arm of the ground reaction forces about the center of mass generates greater fluctuations in angular plane momentum ^81^, which in and of themselves are destabilizing. These changes may be possible because the margins for error of balance control are inherently increased, making wide-base walking more robust to external disturbances. The current results therefore suggest that vestibular contributions may also decrease under walking conditions with reduced control effort to maintain mediolateral stability. This is in line with recent findings in patients with vestibular hypofunction who walk slower, with increased cadence and wider steps ^82-86^. Our results suggest that they may adopt these changes in walking behavior to be less dependent on vestibular input.

The possibility that vestibular contributions are modulated with the control effort (i.e. stabilization demand) of gait is further supported by our narrow-base walking results. Both the dynamic stability of gait and vestibular-evoked ground reaction forces during narrow-base walking increased relative to normal walking. The increase in dynamic stability (i.e. decreasing LDE) in particular is thought to be required to constrain torso motion to margins of error within which walking with narrow steps can be maintained ^34^. Indeed, foot placement during narrow-base walking is more tightly coupled to center-of-mass motion ^30,31^ and participants maintain smaller step-to-step oscillations ^87^. Our results show that under these highly regulated conditions, this increase in control effort may be driven, at least in part, by a net increase in vestibular input (as reflected in the ground reaction forces). This overall increase in vestibular contribution to mediolateral stability, however, does not seem to originate from a general upregulation of vestibular responses in all muscles. Instead, we found decreasing and unchanging vestibular responses in medial gastrocnemius and gluteus medius muscles, respectively, while vestibular responses in erector spinae muscles increased. These muscle specific changes align with the scaling of evoked responses according to the muscle’s involvement in correcting for imposed errors ^11,26^. For example, moments generated by the ankle muscles, such as the medial gastrocnemius, strongly contribute to balance corrections through push-off modulation ^88^. During narrow-base walking, however, push-off modulation is not a viable option since the push off force has only a minimal moment arm. Similarly, regulation of mediolateral foot placement as provided by the gluteus medius ^11,57^ is rendered less effective, since narrow-base walking constrains the foot to a restricted range. Instead, humans more commonly rely on the modulation of the torso’s angular moment to maintain upright balance ^61,87^. Although the muscles around the ankles and hips also contribute to frontal plane angular momentum during normal walking ^81^, the direct influence of spinal muscles on trunk motion may make them more suited to contribute to balance during narrow walking by producing or responding to rapid trunk tilts. A limitation to this interpretation, however, is that we cannot rule out the influence of arm movements, which are known to influence the stability of human gait ^89-91^, since we did not measure arm movements or provide specific instructions to participants to control their movement. Nevertheless, our erector spinae results indicate that spinal muscles provide a key, and perhaps primary, contribution to the net vestibular output to ground reaction forces to maintain upright balance during narrow-base walking.

Significant coherence between the electrical stimulus and erector spinae muscle activity in all conditions also demonstrates that the phasic contribution of this particular axial muscle to whole-body mediolateral stability can be flexibly modulated to address varying stabilizing demands. This is similar to this muscle’s differential response to vestibular disturbances in standing and sitting ^92^. Others have reported, however, that erector spinae muscles, unlike lower limb muscles, maintain a fixed response sensitivity to vestibular input throughout all phases of walking ^93^. This follows a similar argument made in neck muscles where vestibulocollic reflexes are maintained regardless of the requirement to maintain head on trunk balance ^94,95^. By delivering square wave EVS pulses at and slightly after (15% of the gait cycle) the heel strike, Guillaud et al. (2020) argued that the invariant response of the muscle to the electrical stimulation at these two time points indicate a fixed vestibular sensitivity throughout the entire gait cycle ^93^. The improved resolution of the time-varying techniques used here, however, reveals that this muscle does indeed modulate its response to vestibular input throughout locomotion: EVS-EMG coherence peaked at heel strike and dropped to near-zero at approximately 40% of the stride cycle. The restricted time-window of the two stimulation occurrences considered (i.e. at and slightly after heel strike) in the study from Guillaud et al.(2020), however, may have masked any changes in EVS-evoked responses. In addition, the modulation of this muscle’s vestibular sensitivity extends across walking conditions, which nearly doubled during narrow-base walking as compared to normal walking.

A limitation of our study is that while vestibular-evoked responses were quantified throughout the gait cycle, the local divergence exponent remains a mean measure of stability for the entire gait cycle. It may therefore be possible that time-varying changes in gait stability contribute to the phase-dependent vestibular responses seen here. While early studies using phase dependent metrics of gait stability suggested that these measures may hold promise ^96,97^, a recent study from our lab doubted their use, as we found only limited correlation between phase dependent gait stability measures and the probability of falling in a simple dynamic walking model ^98^. However, if sufficient phase dependent gait stability measures become available, the relationship between phase dependent gait stability and phase-dependent vestibular responses may be an interesting study topic.

In conclusion, we have shown that the muscle and whole-body responses evoked by a vestibular stimulation differ according to the gait stabilization demands. When stability is increased by external support, the muscle and whole-body responses to the vestibular stimulus are substantially reduced. During wide-base walking vestibular-evoked muscle and force responses also decrease, though these changes in vestibular contribution are not accompanied by increased dynamic gait stability. Conversely, narrow-base walking produced complex muscle-specific responses that resulted in an increase in the net vestibular contribution to ground reaction forces and increased stability of gait. Overall, our results show that although the vestibular control of gait stability may vary with frontal plane stability, they critically depend on the stabilization demands (i.e. control effort) needed to maintain stable walking patterns.

## Acknowledgements

RMM was funded by CAPES (PDSE 19/2016). SMB was funded by a VIDI grant (016.Vidi.178.014) from the Netherlands Organization for Scientific Research (NWO). PAF received funding from the Netherlands Organization for Scientific Research (NWO #016.Veni.188.049).

## Author contribution statement

JVD and PAF contributed to the conception or design of the work. RMM, SMB and PAF contributed to the acquisition, analysis, or interpretation of data. SMB and PAF made the figures. All authors wrote the main manuscript text, reviewed the manuscript and have approved the submitted version.

